# A Deep Learning Approach for Tissue Spatial Quantification and Genomic Correlations of Histopathological Images

**DOI:** 10.1101/2020.03.10.985887

**Authors:** Zixiao Lu, Xiaohui Zhan, Yi Wu, Jun Cheng, Wei Shao, Dong Ni, Zhi Han, Jie Zhang, Qianjin Feng, Kun Huang

**Author notes:** Co-first authors (Zixiao Lu, Xiaohui Zhan). Corresponding authors (Feng Q), (Huang K).

## Abstract

Epithelial and stromal tissue are components of the tumor microenvironment and play a major role in tumor initiation and progression. Distinguishing stroma from epithelial tissues is critically important for spatial characterization of the tumor microenvironment. We propose an image analysis pipeline based on a Convolutional Neural Network (CNN) model to classify epithelial and stromal regions in whole-slide images. The CNN model was trained using well-annotated breast cancer tissue microarrays and validated with images from The Cancer Genome Atlas (TCGA) project. Our model achieves a classification accuracy of 91.02%, which outperforms other state-of-the-art methods. Using this model, we generated pixel-level epithelial/stromal tissue maps for 1,000 TCGA breast cancer slide images that are paired with gene expression data. We subsequently estimated the epithelial and stromal ratios and performed correlation analysis to model the relationship between gene expression and tissue ratios. Gene Ontology enrichment analyses of genes that were highly correlated with tissue ratios suggest the same tissue was associated with similar biological processes in different breast cancer subtypes, whereas each subtype had its own idiosyncratic biological processes governing the development of these tissues. Taken all together, our approach can lead to new insights in exploring relationships between image-based phenotypes and their underlying genomic data and biological processes for all types of solid tumors.

## Introduction

Most solid tumors are composed of many tissue types including cancer cells, stroma, and epithelium. The interaction of tissues within such complex neoplasms defines the tumor microenvironment and this variably contributes to cancer initiation, progression, and therapeutic responses. For example, breast cancer epithelial cells of the mammary ducts are commonly the site of tumor initiation, while stromal tissue dynamics drive invasion and metastasis [1]. Tumor-to-stroma ratios of H&E stained images are therefore an important prognostic factor [2,3], and distinguishing stromal from epithelial tissue in histological images constitutes a basic, but crucial, task for cancer pathology. Classification methods (*i.e*. pre-processing, training classifiers with carefully selected features, and patch-level classification) are the most common automated computational methods for tissue segmentation [4,5]. For instance, Bunyak et al. [6] combined traditional feature selection methods and classification methods to perform segmentation of epithelial and stromal tissues on a tissue microarray (TMA) database. While this approach is viable, it can be time-consuming and inefficient given the feature selection process. Convolutional Neural Networks (CNN) models have the potential to improve analysis time and performance. Recently, deep CNN models have greatly boosted the performance of natural image analysis techniques such as image classification [7], object detection [8] and semantic segmentation [9,10], and biomedical image analysis [11–13]. Additionally, Ronneberger et al. [14] proposed implementation of a U-Net architecture to capture context and a symmetric expanding path that enables precise localization in biomedical image segmentation. CNN models have also been combined with traditional approaches to enhance the segmentation performance of epithelial and stromal regions [11,12].

Despite breakthroughs in the application of CNN models to medical image analysis, automated classification of epithelial and stromal tissues in Whole Slide Tissue Images (WSI) remains challenging due to the large size of WSI. WSI contain billions of pixels, and machine learning methods are limited by the technical hurdles of working with large datasets [13]. Several solutions based on deep learning for classification of WSI have been proposed. A context-aware stacked CNN was proposed for the classification of breast WSI into multiple categories, such as normal/benign, ductal carcinoma in situ and invasive ductal carcinoma [15]. Saltz et al. presented a patch-based CNN to classify WSI into glioma and non-small-cell lung carcinoma subtypes [16,17].

Additionally, commercial software has been developed to aid in quantitative and objective analyses of tissue WSI. Among them is GENIE (Leica/Aperio), a tool with proprietary algorithms which incorporate deep learning. While many of its functionalities are designed to handle specific biomarkers using immunohistochemical (IHC) or fluorescent images, for H&E images, tissue segmentation requires user-defined regions of interests (ROI). Similarly, HALO (Indica Labs) and Visiopharm (Hoersholm) provide a toolbox for histopathological image analysis. The toolbox includes unsupervised algorithms for tissue segmentation that require manual configuration of parameters and usually underperform supervised methods. The AQUA system (HistoRx) focuses on estimating tissue scores on TMA based on IHC staining by measuring protein expression within defined ROI. Therefore, reliable systems that enable both fully-automatic tissue segmentation and quantified analysis for H&E whole-slide images are still in great demand.

In this work, we propose a WSI processing pipeline that utilizes deep learning to perform automatic segmentation and quantification of epithelial and stromal tissues for breast cancer WSI from The Cancer Genome Atlas (TCGA). The TCGA data portal provides both clinical information and paired molecular data [18,19]. This offers the opportunity to identify relationships between computational histopathologic image features and the corresponding genomic information, which greatly informs research into the molecular basis of tumor cell and tissue morphology [20–22], as well as important issues such as immune-oncology therapy [17].

We first trained and validated a deep CNN model on annotated H&E stained histologic image patches, then successfully applied the WSI processing pipeline to process 1,000 TCGA breast cancer WSI to segment and quantify epithelial and stromal tissues. Spatial quantification and correlations with genomic data of both tissue types for three breast cancer subtypes (ER-positive, ER-negative and triple negative) were estimated based on the high-resolution global tissue segmentation maps. Gene Ontology (GO) enrichment can indicate when such tissues are associated with similar biological processes in different breast cancer subtypes, whereas each subtype has its own idiosyncratic biological processes governing the development of these tissues. These results are consistent with underlying biological processes for cancer development, which further affirms the robustness of our image processing method.

Spatial characterization of different tissues in histopathological images has shown significant diagnostic and prognostic value, but human assessment of these features is time-consuming and often infeasible for large-scale studies. This study contributes an innovative automated deep-learning analysis pipeline that will enable rapid, accurate quantification of epithelial and stromal tissues from WSI of cancer samples. Such approaches are useful because they may be used for the quantification of tissue-level epithelial/stromal/cancer phenotypes, which in turn may be integrated with other biomedical data. For this reason, we demonstrate how model-generated outputs may be correlated with gene expression and how this may lead to new insights about genetic mechanisms that contribute to tumor microenvironment variability in breast cancer. Additional contributions of this manuscript are that the approach, data, and demonstrated use of the pipeline could be applied to other cancers to improve tissue quantification. To the best of our knowledge, this is the first study to provide pixel-level tissue segmentation maps of TCGA image data.

## Method

### Datasets

Two breast cancer image sets were used in this study: (1) The Cancer Genome Atlas (TCGA) portal; (2) the Stanford Tissue Microarray Database (sTMA) [2]. The sTMA database consisted of a total of 157 H&E stained rectangular image regions (1128 × 720 pixels) using 20X objective lens, which were acquired from two independent cohorts: 106 samples from Netherlands Cancer Institute (NKI) and 51 samples from Vancouver General Hospital (VGH). Each image of sTMA was manually annotated with epithelial and stromal tissues by pathologists. The TCGA cohort samples include matched H&E stained WSI, gene expression data, and clinical information. Patients with missing expression data or images with cryo-artifacts deemed too severe were excluded, leaving a selected set of 1,000 samples. Since the TCGA clinical information includes subtyping information, we further categorized the selected samples into three breast cancer subtypes for more specific biological analysis: ER-positive, ER-negative and triple negative. Demographic and clinical information for both sTMA and TCGA cohorts are summarized in **Table 1**.

**Table 1.** Demographic and clinical characteristics.

| Cohort | SubGroup | Image Type | Image Number | Total |
| --- | --- | --- | --- | --- |
| TMA | NKI | H&E stained image<br>region (1128 * 720) | 106 | 157 |
|  | VGH |  | 51 |  |
| TCGA | ER-positive | Whole-slide image | 773 | 1000 |
|  | ER-negative |  | 227 |  |
|  | Triple negative |  | 112 |  |

*Note:* For TCGA cohort, samples in Triple Negative subgroup also belong to ER-negative subgroup.

### Overview of the workflow

**Figure 1** outlines our workflow for both image processing and biological analysis. **Figure 1A** shows the detailed structure of our deep CNN model for tissue segmentation. **Figure 1B** is the whole-slide image processing pipeline. **Figure 1C** shows an overview of the biological analysis of gene expression data and image features. Details of each part are described in the following subsection.

**Figure 1.**
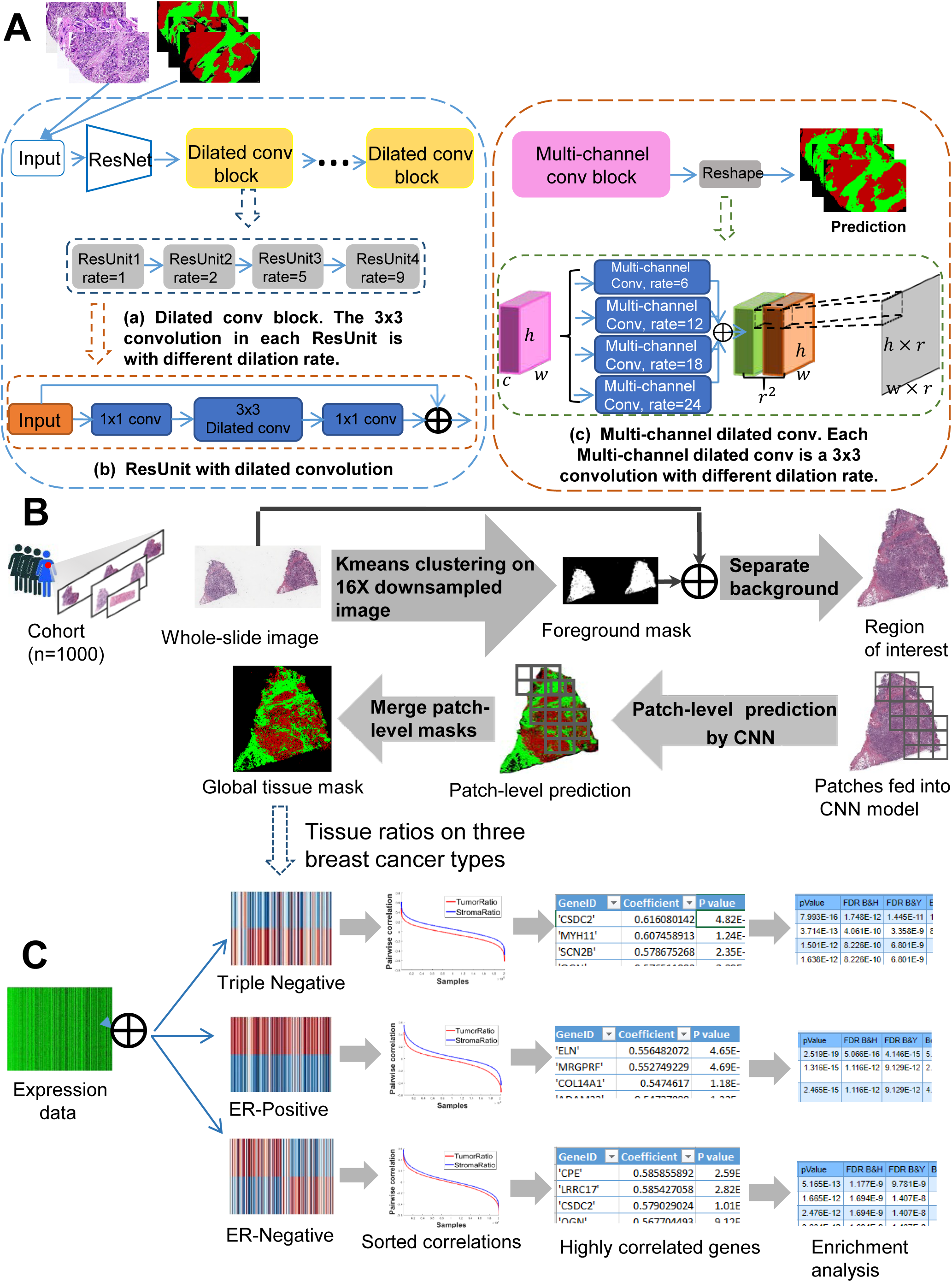
Workflow for image processing and biological analysis. **A**. Detailed structure of our deep CNN model for segmentation. **B**. Whole-slide image processing pipeline. **C**. Overview of biological analysis of gene expression data and image features.

### CNN model for tissue segmentation

Given an RGB image of height *H*, width *W*, and color channels *C*, the goal of segmentation is to predict a label map with size *H* × *W* where each pixel is labeled with a category. CNN-based framework for segmentation fundamentally consists of encoding and decoding counterparts.

The encoding block is derived from classification models which perform down-sampling operators to capture global information from input images. Max-pooling is the most commonly adopted operations in encoding, which integrates neighbouring pixels to learn invariance from local image transformation. More recently, dilated convolution was proposed to control spatial resolution, and thus enable dense feature extraction. Given a 1-D input signal *x[i]* with a filter *w[k]* of length K, the output of dilated convolution is defined as:

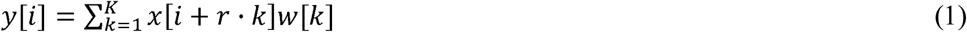

where *r* is the stride in the sampling input signal, referred to as *rate*. By filling zeros between pixels in the filter, dilated convolution can enlarge receptive fields without substantially increasing computational cost.

We carefully constructed our deep hierarchical segmentation model using specific strategies in both encoder and decoder, as shown in **Figure 1A**. The ResNet-101 structure [7], which contains 101 convolution layers, was adopted as the backbone of our proposed model. Since dilated convolution inserts zeros between pixels in the filter, it can enlarge receptive fields without substantially increasing computational cost. The encoder of our model inherited the first three blocks of ResNet-101, while the rest were modified into six dilated convolution blocks, each of which further contained four ResUnits with different dilation rates. This configuration was inspired by the success of the atrous spatial pyramid pooling (DeepLab-ASPP) approach from Chen et al. [10], which captures objects as well as image context at multiple scales, and thus robustly improves the segmentation performance. In our work, the modification of convolution layers was conducted to ensure that our encoder learned both tissue structures and contextual information for the next phase of processing. In the decoding step, we adopted a multi-channel convolution approach to generate high-resolution segmentation maps. Given a feature map of dimension *h × w × c*, multi-channel convolution first generated features of *h × w ×* (*r*^2^ × *c*), where *r* is the upsampling rate. Then the features were reshaped to obtain upsampled features of *H′ × W′ × c, where H′ = h × r, W′ =w × r*. To this end, we stretched each individual pixel in the small feature map to the channel of *r*^2^ × *c* so that it corresponded to a fixed area (*r* × *r*) in the upsampled output map. We applied four parallel dilated multi-channel convolutions with a range of dilation rates and added all of their outputs pixel by pixel in order to further exploit multi-scale contextual information from the encoding feature map.

We next used sTMA to train our CNN model in a five-folder-cross-validation. The proposed model was implemented using MXNet toolbox. Parameters in the encoder were initialized with pre-trained weights from Deep-Lab V2 [10], while the decoder layers were randomly initialized by Xavier method. Due to GPU memory limitations (8 GB for GeForce GTX 1080), we randomly cropped 600 × 600 patches from the raw images and performed random mirror and random crop as data augmentation in the training stage.

### WSI processing pipeline

During biopsy slide examination, pathologists search for a region-of-interest (ROI) that contains cancer cells and conduct diagnostic assessment. Inspired by these human analysis steps, we built an automatic pipeline to perform tissue segmentation on WSI, as shown in **Figure 1B**. Our WSI processing pipeline consists of two parts: 1) automatic identification of ROI, and 2) epithelial and stromal tissue segmentation on the ROI. Given a WSI *I*, we first downsampled *I* into *I*′ at a factor of 16 in both horizontal and vertical directions. Then we converted *I*^′^ from RGB color space to CIELAB color space (*L*\**a*\**b*\*), denoted as 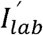. Since the *L*\* channel in *L*\**a*\**b*\* color space represents the brightness, we extracted the *a*^*^ and *b*^*^ values representing color components in 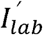 and obtained a new image 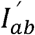. Each pixel in 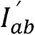 is then a 2-dimentional vector. Next, we applied K-means clustering algorithm (K=2) to divide the pixels of 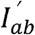 into two groups. Considering that corners of pathology images are usually unstained, we classified pixels in the same cluster as the upper-left pixel in 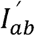 as background, while the other pixels were classified as foreground. In this way, we generated a binary mask *M*^1^, where 0 and 1 in *M*^1^ correspond to background and foreground pixels in 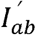, respectively. Denoting the smallest rectangle region that contains the largest connected component in *M*^1^ as *F_m_*, we identified the ROI *F_I_* by mapping the coordinates of *F_m_* onto *I*. Finally, *F_I_* was cropped from *I* for downstream processing.

We split *F_I_* into patches of 1128 × 720 pixels to fully utilize the proposed CNN model for tissue segmentation. Patches with more than 80% background were discarded. The retained patches were then fed into the CNN model and all the patch-level predictions were combined to generate a global tissue mask *M*^2^ for *F_I_*.

### Tissue quantification and biological analysis

We applied our WSI processing pipeline on 1,000 TCGA breast cancer WSI for further biological analysis, as shown in **Figure 1C**. For each WSI *I*, we performed tissue spatial quantification based on its tissue mask *M*^2^ derived from our method. The two tissue ratios, *Ratio_epi_* and *Ratio_stro_*, that characterize the ratio of epithelial tissue areas and stromal tissue areas to overall tissue areas were estimated as:

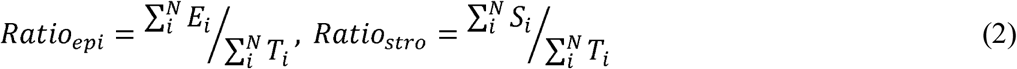

where *T*_*i*_, *E*_*i*_ and *s*_*i*_ represent the number of pixels classified as foreground, epithelial and stromal in the ith valid patch in F_*I*_ respectively, and *N* represents the total number of valid patches in *F*_*I*_.

To explore the relationships between gene expression data and tissue ratios in different breast cancer subtypes, we divided all TCGA samples into three types: ER-positive, ER-negative, and triple negative, as seen in **Table 1**. Then, we computed the Spearman correlation coefficients between gene expression data and the two tissue ratios *Ratio*_*epi*_ and *Ratio*_*stro*_ for each breast cancer subtype. Next, we sorted all the Spearman coefficients and selected the gene symbols which were in the top 1% of correlation coefficients with *Ratio*_*epi*_ and *Ratio*_*stro*_ for each breast cancer subtype. For the selected gene symbols, we performed Gene Ontology (GO) enrichment analysis on them using WebGestalt [23]. Meanwhile, the Overrepresentation Enrichment Analysis (ORA) with Bonferroni adjustment methods was also used to determine statistical significance of the enrichment. Genes presented by the “Genome” platform were used as the reference gene. Finally, the top 10 enriched biological process categories were selected to reveal the biological process underlying the development of epithelial and stromal tissues for each breast cancer subtype.

## Results

### Validation of CNN model

We evaluated the effectiveness of our proposed deep CNN model on segmentation of epithelial and stromal tissues by testing and comparing our model with several state-of-the-art methods [11,12,24,25]. Our model outperformed all of these methods based on a comparison of classification accuracies and achieved an average accuracy of 91.02% on the whole sTMA dataset (NKI + VGH), as shown in **Table 2** and **Table 3**. Visual segmentation results also demonstrated that our model could accurately classify epithelial and stromal tissues (**Figure 2**). Note that in the ground truth data, some areas belonging to epithelia have been overlooked and incorrectly annotated as background (an example is shown in the third row of **Figure 2**). However, our model still yielded correct predictions on this area (marked by a black circle in **Figure 2**). This indicates that our model is robust enough to make the right judgment, even under misleading supervision. We believe this is valuable for future work in biomedical image tasks with only partial or inaccurate annotations.

**Table 2.** Evaluation of CNN model on NKI and VGH.

| Datasets | Models | TPR | TNR | PPV | NPV | FPR | FDR | FNR | ACC | F1 | MCC |
| --- | --- | --- | --- | --- | --- | --- | --- | --- | --- | --- | --- |
| **NKI** | Xu.et [12] | 86.31 | 82.15 | 84.11 | 84.60 | 17.85 | 15.89 | 13.66 | 84.34 | 85.21 | 68.60 |
|  | CNN only [11] | 81.34 | 82.89 | 84.11 | 80.05 | 17.11 | 15.89 | 18.57 | 81.69 | 82.75 | 64.24 |
|  | CNN+HFCM [11] | 89.48 | 85.96 | 85.94 | 89.50 | 14.04 | 14.06 | 10.52 | 87.19 | 87.68 | 75.44 |
|  | Our model | **90.71** | **89.83** | **90.81** | **89.72** | **10.17** | **9.19** | **9.29** | **90.29** | **90.76** | **80.54** |
| **VGH** | Xu.et [12] | 88.29 | 88.40 | 89.93 | 86.55 | 11.60 | 10.07 | 11.71 | 88.34 | 89.10 | 76.59 |
|  | CNN only [11] | 90.32 | 88.15 | 92.98 | 83.97 | 11.85 | 7.02 | 9.68 | 89.14 | 91.63 | 77.70 |
|  | CNN+HFCM [11] | **91.96** | **92.21** | **95.45** | 86.59 | **7.79** | **4.55** | **8.04** | 91.04 | **93.67** | **83.10** |
|  | Our model | 91.37 | 91.49 | 92.37 | **90.38** | 8.51 | 7.63 | 8.63 | **91.42** | 91.87 | 82.80 |

*Note:* From the third to the last column are the ten evaluations metrics. Value in bold represents the best result under each metric among different models.
\*TPR (True positive rate) = $TP / (TP + FN)$ ; TNR (True negative rate) = $TN / (FP +$ $TN)$ ; PPV (Positive Predictive Value) = $TP / (TP + FP)$ ; NPV (Negative Predictive
Value) = $TN / (FN + TN)$ ; FPR (False positive rate) = $FP / (FP + TN)$ ; FDR (False
Discovery Rate) = $1 - TP / (TP + FP)$ ; FNR(False Negative Rate) = $FN / (FN + TP)$ ;
ACC (Accuracy) = $(TP + TN) / (TP + FP + TN + FN)$ ; F1\_score = $2 * TP / (2 * TP + FP$ $+ FN)$ ; MCC (Matthews Correlation Coefficient) = $(TP * TN - FP * FN) /$ $\sqrt{(TP + FP) * (TP + FN) * (TN + FP) * (TN + FN)}$ . TP, FP, TN and FN represent the true positive, false positive, true negative and false negative, respectively.

**Table 3.**
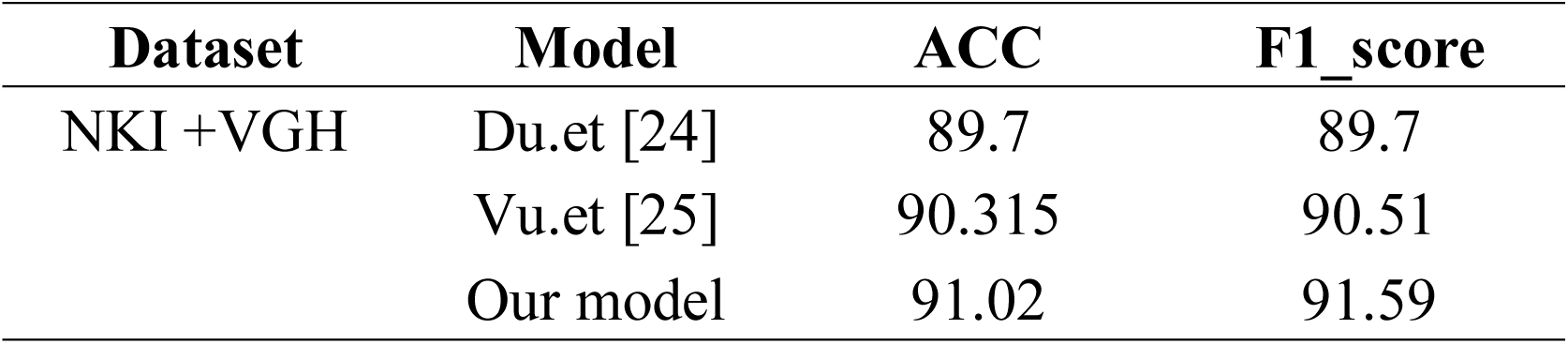
Quantitative evaluation on the whole TMA dataset.

| Dataset | Model | ACC | F1_score |
| --- | --- | --- | --- |
| NKI +VGH | Du.et [24] | 89.7 | 89.7 |
|  | Vu.et [25] | 90.315 | 90.51 |
|  | Our model | 91.02 | 91.59 |

**Figure 2.**
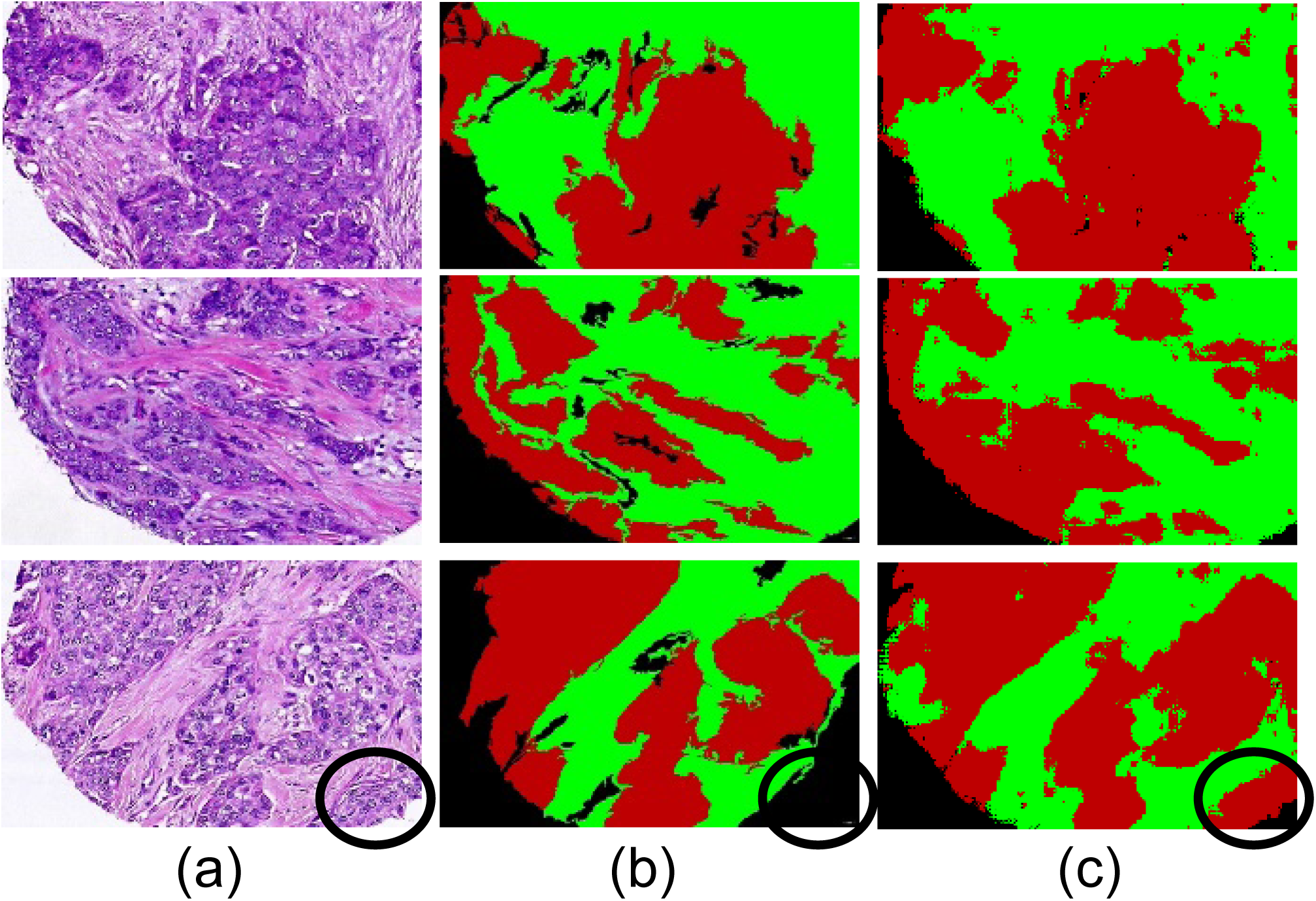
Segmentation results on TMA. Column (a) are raw images; column (b) are annotations by pathologists; column (c) are predictions of the proposed model. Red, green and black areas in column (b) and (c) represent epithelial, stromal and background regions in raw images, respectively. Note that in the last row, the overlooked tumor area (Marked with black circle) is still well recognized by our model.

### Tissue segmentation and quantification on WSI

We validated the trained CNN model on 171 image patches each from the TCGA breast cancer slide images annotated with epithelial/stromal tissues by two domain experts. The validation results indicated that our model was robust enough to predict credible tissue mask for the TCGA dataset (Table T1 and Figure S1). We then applied the trained CNN model to the tissue segmentation of 1,000 whole-slide images from three TCGA breast cancer subtypes. Visual results showed that our pipeline could robustly identify epithelial/stromal tissues in whole-slide images (**Figure 3)**.

**Figure 3.**
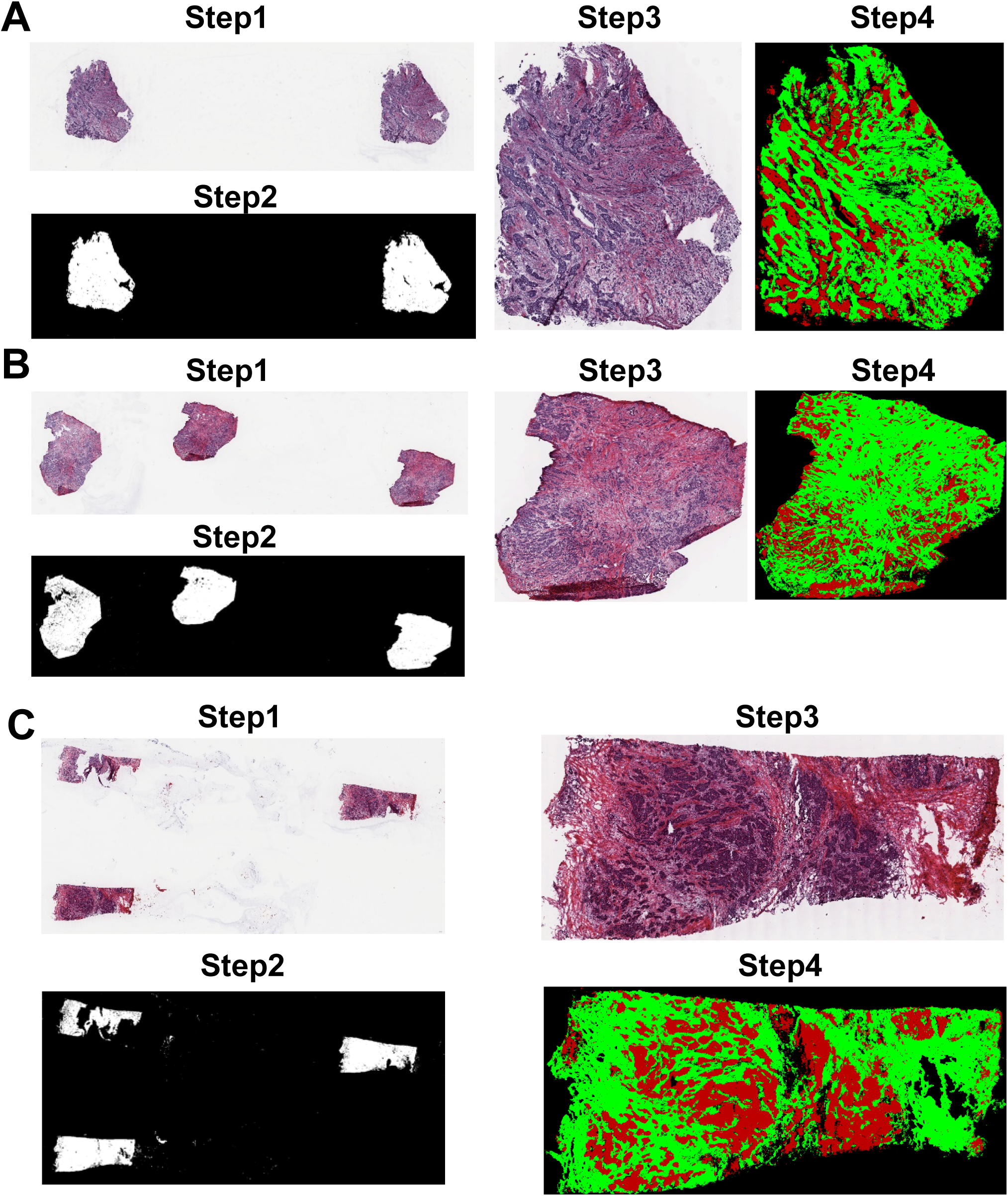
Segmentation results on TCGA WSIs. For each TCGA whole-slide image **A, B, C: Step 1** represents the WSI; **Step 2** represents the background map of WSI; **Step 3** represents the region of interest (ROI) in the WSI of raw resolution; **Step 4** represents the tissue segmentation result of ROI. Red, green and black areas in **Step 4** represent the predicted epithelial, stromal and background regions, respectively.

Ratios of epithelial and stromal tissue areas to overall tissue areas were estimated based on the WSI segmentation results. Wide differences in tissue ratios were seen among different breast cancer subtypes (**Figure 4**). ER-positive images were predominantly enriched with stromal tissues with a mean stromal ratio of 72.8%, while triple negative images were abundant in epithelial tissues with a mean epithelial ratio of 63.56%. Epithelial and stromal tissues were nearly equivalent for ER-negative images with mean ratios of 49.35% and 50.65%, respectively.

**Figure 4.**
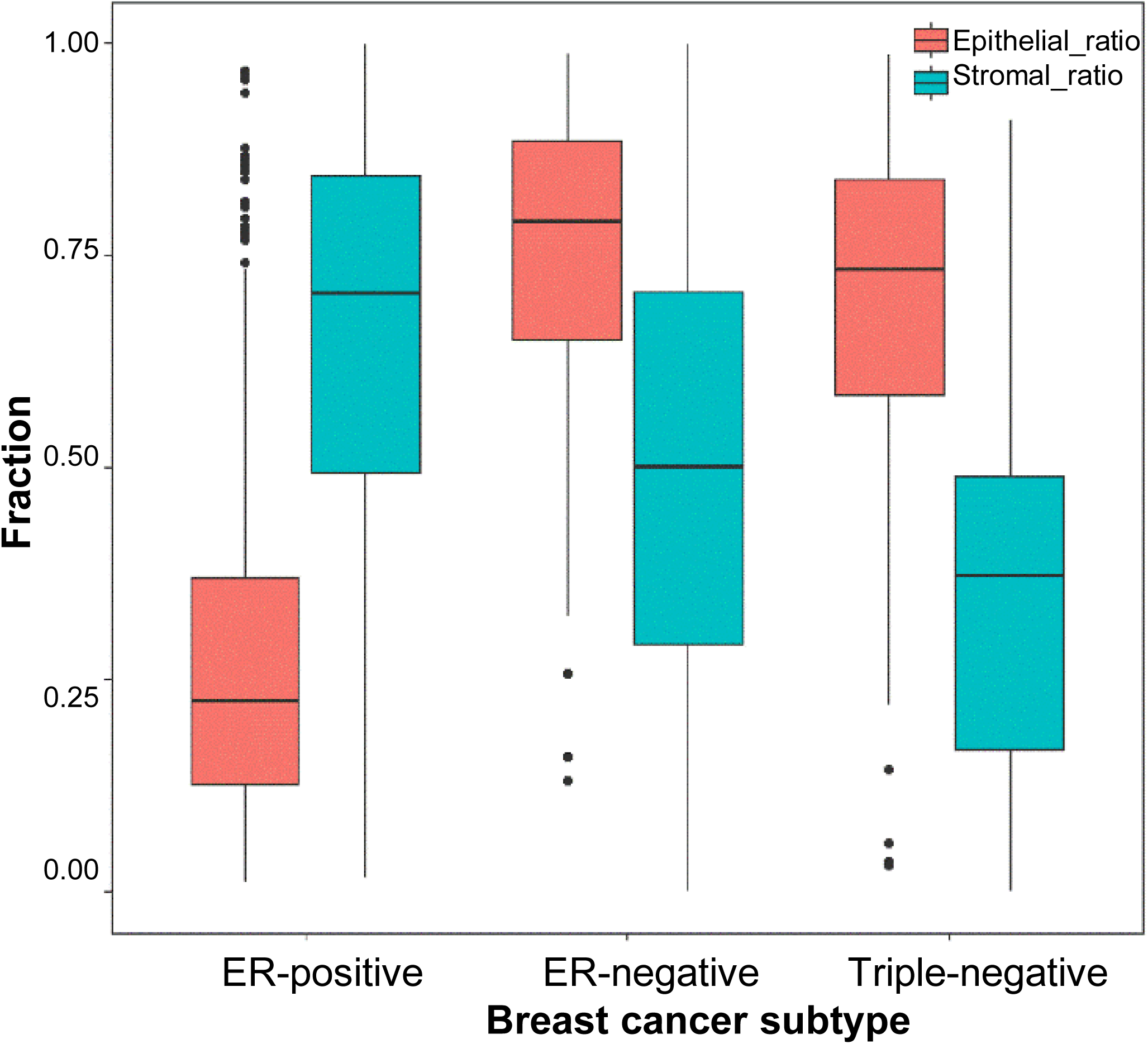
Tissue distribution on different breast cancer subtypes. The Variable Epithelial_ratio, Stromal_ratio represent the ratios of epithelial tissue areas and stromal tissue areas to overall tissue areas, respectively.

### Tissue-specific functional analysis

We explored which genes contributed to the development of different tissues in various subtypes of breast cancers by computing pairwise Spearman correlation coefficients between gene expression data and both tissue ratios. Genes in the top 1% of correlation with tissue ratios in each subtype of breast cancer were selected for further analysis. We then performed functional Gene Ontology (GO) analysis for the selected gene-sets. Genes correlated with the epithelial tissues were enriched in biological processes during the cell cycle, among which sister chromatid segregation, nuclear division, and mitotic cell cycle are the most commonly enriched GO terms shared by the three breast cancer subtypes. However, we also observed specifically enriched GO terms and genes for each subtype that correspond to different cell cycle stages. The Growth phase related genes including G1 phase and G2 phase were specifically enriched for the ER-positive subtype, whereas Mitotic phase genes were specifically enriched for the triple negative subtype, and S phase related genes were specific for the ER-negative subtype.

Similarly, such patterns of shared high-level biological processes with specific functions were also observed for the stromal tissues. For the stromal tissue, the most significantly enriched GO biological process terms were all related to the development of the tumor microenvironment, including vasculature development, cellular component movement, and growth factor stimuli-related GO functions which were shared among the three breast cancer subtypes. For the ER-positive subtype, angiogenesis-related genes were specifically enriched, while for the triple negative subtype, muscle structure genes (especially the ones related to actin fibers and cytoskeleton) were specifically enriched. In addition, for the ER-negative subtype, growth factor genes were enriched. Altogether, our results (**Figure 5**) suggest that even though the same tissue was associated with similar biological processes in different subtypes, each subtype still had its idiosyncratic biological processes governing the development of these tissues.

**Figure 5.**
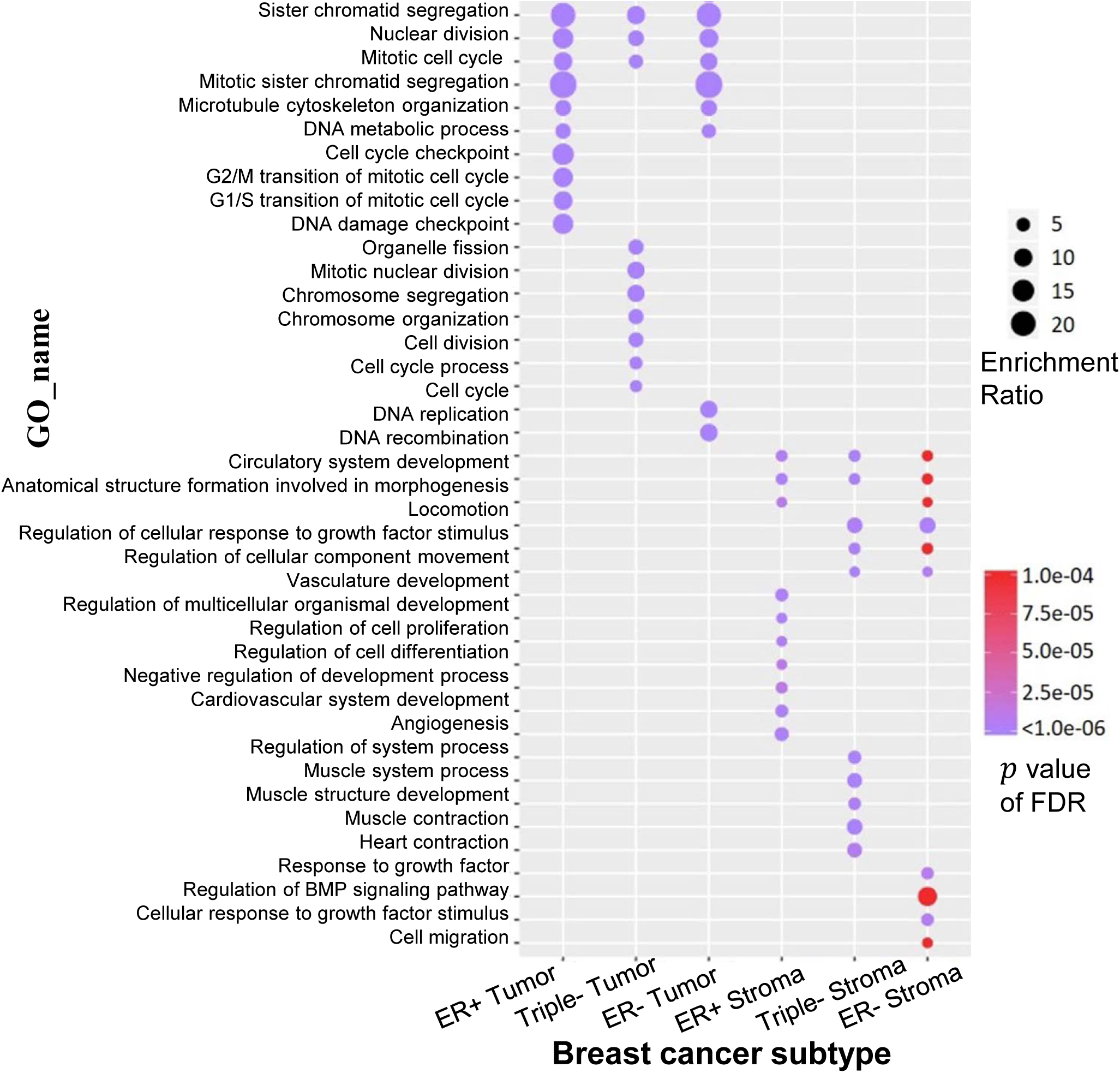
Results of GO enrichment analysis. Dots represent most significantly enriched Biological Process term for each cancer subtype with color coding: purple indicates high enrichment, red indicates low enrichment. Sizes of dots represent the ratio of enrichment (GO category). FDR is the method used for multiple comparison correction.

### Other applications

Our WSI processing pipeline can be easily applied to histological images of other types of cancers. The global tissue segmentation maps we have presented could also be used for other more specific computational analysis. For example, global morphological features of different tissues could be estimated for better survival prediction [22,26], and lymphocytes in different tissues could be distinguished for observation of more detailed immune response. Imaging data resources have not been exploited to the degree of the other TCGA molecular and clinical outcome resources, likely because automatic image annotation is still impeded by data volume challenges. In this manuscript we presented global tissue maps of all the TCGA breast cancer WSI, and it is our aspiration that they will facilitate further exploration and utilization of these imaging data for various cancers.

## Conclusions

Epithelial and stromal regions of tumors, as well as their spatial characterizations in histopathology images, play a very important role in cancer diagnosis, prognosis, and treatment. Recently, some research studies have focused on developing systems for automatically analyzing H&E stained histological images from tissue microarrays in order to predict prognosis [26,27]. In contrast, our approach is aimed at whole slide images (WSI) rather than manually extracted regions since WSI provide much more comprehensive information, including heterogeneity. Mackie et al. [28] summarized the research progress and challenges facing the application of big data quantitative imaging to cancer treatment, focusing on 3D imaging modalities including CT, PET, and MRI. Our quantitative analysis of histopathology images complements and extends this work in terms of data modality and size, application areas, and computational challenges.

Based on our global tissue quantification, distinct differences were observed in the enriched GO terms for epithelial and stromal tissues [29]. At the same time, highly overlapping biological properties were observed in the same tissue across different subtypes, all tied to cancer progression in one way or another. For example, in epithelial tissue, genes from cell cycle-related processes were significantly enriched. Previous studies have addressed that sustaining proliferative signaling is one of the hallmarks of cancer, during which cell cycle plays quite an important role [30]. In addition, *CDK4/6* inhibitors (such as Palbociclib and ribociclib) target this biological process [31,32]. For stromal tissue, genes related to the tumor microenvironment were significantly enriched (e.g., vasculature and locomotion). Vasculature is vital for inducing angiogenesis, which is another important hallmark of cancer.

Additionally, we observed differences in biological processes between different subtypes resulting from tumor heterogeneity. Specific biological process features for each subtype were also identified among the same tissue. For epithelial tissue, genes associated with different stages of the cell cycle were specifically enriched for different subtype. For ER-positive breast epithelia, we found that G1 and G2 phase-related GO terms were enriched, among which G2/M transition is an important element. Wang et al. [27] have highlighted the importance of G2/M transition in ER-positive breast cancer. For the triple negative subtype, we found that M phase related GO terms were enriched, during which chromosome segregation plays a key role. Witkiewicet et al. [33] have shown the close relationship between chromosome segregation (*PLK1*) with triple negative Breast Cancer. Similarly, angiogenesis related biological processes were significantly associated with the stroma of the ER-positive subtype. Previous studies have indicated that vasculature is one of the important components for tumor stroma [34], as stromal cells can build blood vessels to supply oxygen and nutrients [35].

While the correlation analysis of this study reveals clear pairwise relationships between morphological and genomic features, there are two major limitations to our approach. First, correlation cannot reveal highly nonlinear relationships or multivariate complication relationships. For instance, Wang et al. [36] demonstrated that complicated morphological features might need to be modeled using multiple genomic features, implying contributions from multiple genetic factors. Similarly, with our data, more sophisticated analysis such as nonlinear correlation analysis can be applied to reveal deeper relationships. Secondly, correlation is not causation. The genes that are strongly correlated with the stromal or epithelial content may not be the underlying driver genes for the development of the tissues. Identification of such key genes requires further incorporation of biological knowledge, as well as future experimental validation.

In summary, our framework provides not only fully automatic and detailed analysis for large H&E stained images based on a state-of-the-art deep learning model, but also integrated analysis of image features and molecular data. The proposed framework enables us to effectively explore the underlying relationships between gene expression and tissue quantification, free from the extensive labelling and annotation that is laborious even to skilled pathologists.

The details about code and data in this manuscript is provided on Github with the link at *https://github.com/Serian1992/ImgBio*.

## Authors” contributions

LZ carried out the pathology image processing, participated in the genetic studies and drafted the manuscript. ZX carried out the enrichment analysis and helped to draft the manuscript. WY participated in the development of methodology. CJ participated in the acquisition of data. SW participated in the development of methodology. HZ participated in the acquisition of data and the development of methodology. ZJ and DN participated in the review and revision of the manuscript. FQ participated in the development of methodology and helped to review and revise the manuscript. HK conceived of the study, and participated in its design and coordination, reviewed and edited the manuscript. All authors read and approved the final manuscript.

## Competing interests

The authors have declared no competing interests.

## Acknowledgements

This work was supported by Indiana University Precision Health Initiative to HK and ZJ, the NSFC-Guangdong United Found of China (No. U1501256) to FQ, and Shenzhen Peacock Plan (No. KQTD2016053112051497) to ZX and DN. We thank Dr. Natalie Lambert, Dr. Bryan Helm and Ms. Megan Metzger for their tremendous help in the discussion and editing of the manuscript.

## Supplementary material

**Figure S1 Qualitative segmentation results on TCGA dataset**

The first column are the raw TCGA images; the second column are annotations by pathologists; the third column are predictions of the proposed model. Red, green and black areas in the annotations and predictions represent epithelial, stromal and background regions in raw images, respectively.

**Table T1 Quantitative evaluation on TCGA dataset**

* TPR (True positive rate) = TP / (TP + FN); TNR (True negative rate) = TN / (FP + TN); FPR (False positive rate) = FP / (FP + TN); FNR(False Negative Rate) = FN / (FN + TP); ACC (Accuracy) = (TP + TN) / (TP + FP + TN + FN); F1_score = 2*TP / (2*TP + FP + FN). TP, FP, TN and FN represent the true positive, false positive, true negative and false negative, respectively.

