## Supplementary figures and images for "A Deep Learning Approach for Tissue Spatial Quantification and Genomic Correlations of Histopathological Images"

### Supplemental Figure 1

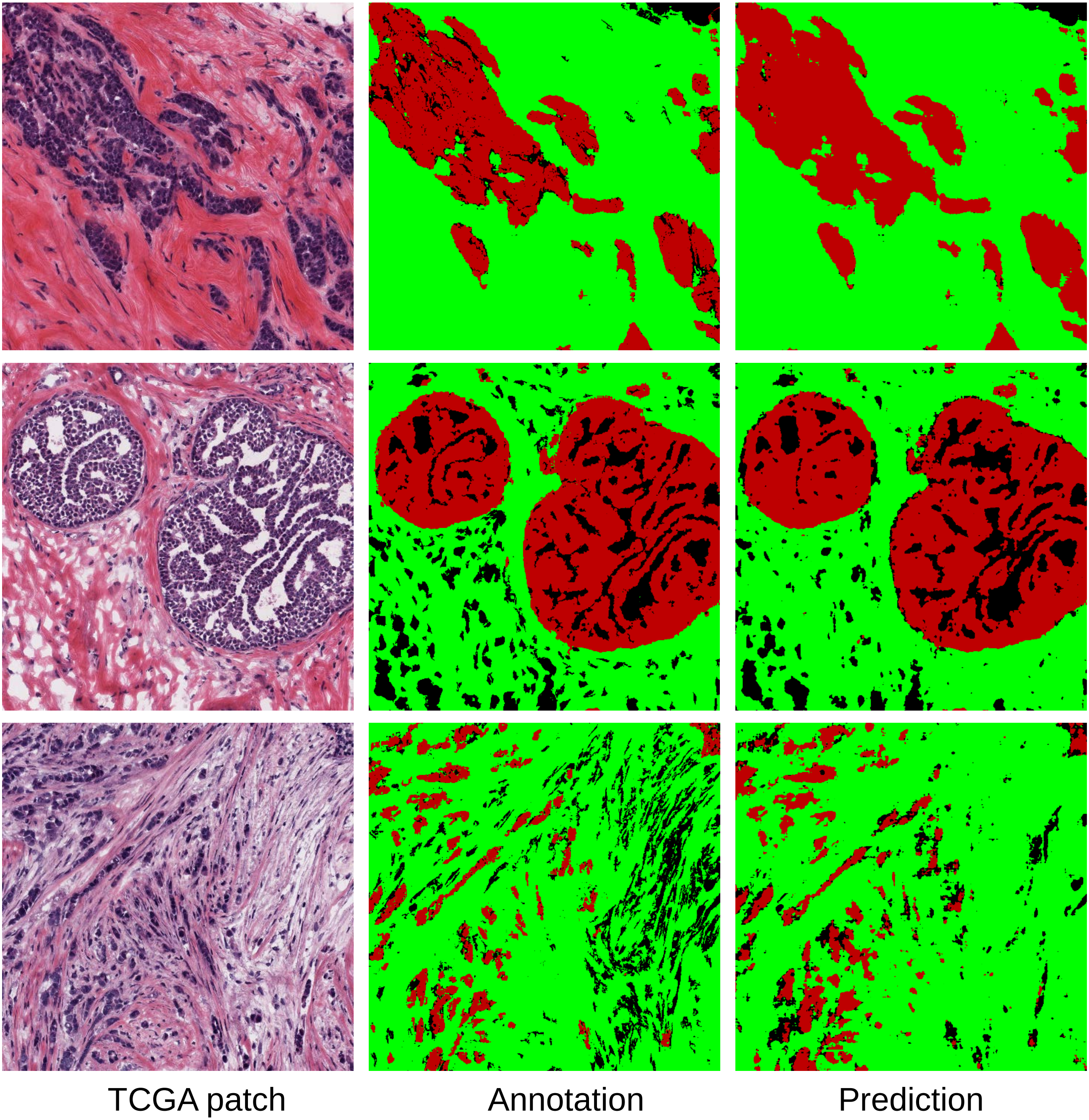
