## Supplemental Table 1 for "A Deep Learning Approach for Tissue Spatial Quantification and Genomic Correlations of Histopathological Images"

**Table T1 Quantitative evaluation on TCGA dataset**

| **Dataset** | **Model** | **TPR** | **TNR** | **FPR** | **FNR** | **ACC** | **F1_score** |
| --- | --- | --- | --- | --- | --- | --- | --- |
| TCGA | Our model | 87.19 | 95.08 | 4.92 | 12.81 | 92.05 | 89.39 |

* TPR (True positive rate) = TP / (TP + FN); TNR (True negative rate) = TN / (FP + TN); FPR (False positive rate) = FP / (FP + TN); FNR(False Negative Rate) = FN / (FN + TP); ACC (Accuracy) = (TP + TN) / (TP + FP + TN + FN); F1_score = 2$*$TP / (2$*$TP + FP + FN). TP, FP, TN and FN represent the true positive, false positive, true negative and false negative, respectively.
